# Anti-stress effects of the glucagon-like peptide-1 receptor agonist liraglutide in zebrafish

**DOI:** 10.1101/2021.01.08.425989

**Authors:** Pablo R. Bertelli, Ricieri Mocelin, Matheus Marcon, Adrieli Sachett, Rosane Gomez, Adriane R. Rosa, Ana P. Herrmann, Angelo Piato

**Author notes:** Correspondence to: Angelo Piato, Ph.D. Programa de Pós-graduação em Ciências Biológicas: Farmacologia e Terapêutica, Instituto de Ciências Básicas da Saúde, Universidade Federal do Rio Grande do Sul (UFRGS), Avenida Sarmento Leite, 500/305, Porto Alegre, RS 90050-170, Brazil. The authors contributed equally to this manuscript (Pablo R. Bertelli, Ricieri Mocelin).

## Abstract

Stress-related disorders are extremely harmful and cause significant impacts on the individual and society. Despite the limited evidence regarding glucagon-like peptide-1 receptor (GLP-1R) and mental disorders, a few clinical and preclinical studies suggest that modulating this system could improve symptoms of stress-related disorders. This study aimed to investigate the effects of liraglutide, a GLP-1R agonist, on neurobehavioral phenotypes and brain oxidative status in adult zebrafish. Acute liraglutide promoted anxiolytic-like effects in the light/dark test, while chronic treatment blocked the impact of unpredictable chronic stress on behavioral and physiological parameters. Taken together, our study demonstrates that liraglutide is active on zebrafish brain and may counteract some of the effects induced by stress. More studies are warranted to further elucidate the potential of GLP-1R agonists for the management of brain disorders.

## 1. INTRODUCTION

Glucagon-like peptide-1 (GLP-1) is an incretin hormone that plays an essential role in regulating blood glucose levels and insulin sensitivity. It is secreted by L-cells in the gut epithelium in response to nutrients, promoting glucose homeostasis (KAHN; HULL; UTZSCHNEIDER, 2006). GLP-1 exerts its physiological actions by binding to the GLP-1 receptor (GLP-1R), which is expressed in abundance in peripheral and central tissues, enabling metabolic, endocrine, behavioral, and cardiovascular regulation in response to homeostatic signals (HOLT; TRAPP, 2016; KRASNER et al., 2014; RUSSELL-JONES, 2009). Studies have also shown the production of GLP-1 in the brain of mammals, where it may play a role in stress regulation by modulating the hypothalamic-pituitary-adrenal (HPA) axis and the autonomic nervous system (LÓPEZ-FERRERAS et al., 2018; ULRICH-LAI; HERMAN, 2009).

GLP-1R agonists are a new drug class developed for glycemic control in diabetic patients. The GLP-1R agonists currently approved for type-2 diabetes mellitus (T2DM) include albiglutide, dulaglutide, exenatide, liraglutide, lixisenatide, and semaglutide. Liraglutide has 97% homology to the human GLP-1 and was approved by the FDA in 2010. Like other GLP-1R agonists, it stimulates insulin secretion and inhibits glucagon release by the pancreas, delays gastric emptying, and reduces appetite (LIM et al., 2009; LÓPEZ-FERRERAS et al., 2018; MOJSOV, 2000; WHICHER et al., 2019). Due to its weight loss properties, liraglutide was repurposed by the Food and Drug Administration (FDA) in 2014 as a treatment for obesity.

The presence of GLP-1Rs in the brain prompted studies on the central effects of GLP-1R agonists, which revealed improvement in cognitive functions, protection against oxidative stress, neuroinflammation, increased synaptic plasticity, up-regulation of brain-derived neurotrophic factor (BDNF), and reduction in apoptosis signaling molecules in rodents (CITRARO et al., 2019; HAN et al., 2020; KRASS et al., 2015; LIU et al., 2019; TAUCHI et al., 2008; WEINA et al., 2018; YAN et al., 2019; ZHANG et al., 2020, 2009, 2019). GLP-1R agonists were also tested in preclinical models of anxiety and depression, including unpredictable chronic stress (UCS) and corticosterone exposure. These studies demonstrated the effects of GLP-1R agonists on forced swimming, tail suspension, elevated plus maze, open field, and light-dark tests, showing antidepressant/anxiolytic proprieties, as well as antioxidant effects in rodents (ABDELAZIZ et al., 2019; CHAVES FILHO et al., 2020; DE SOUZA et al., 2019; HAN et al., 2020; KRASS et al., 2015; LIU et al., 2019; WEINA et al., 2018). However, the effects of GLP-1R agonists and their potential mechanism of action in mental disorders stress-related in zebrafish remains unknown.

Zebrafish has been progressively used in pharmacological and neuroscience studies because of the high similarities in physiology and metabolism compared to mammals. Despite the functional differences between fish and mammalian GLP-1s regarding glucose metabolism, they seem to be interchangeable in their functions (MOMMSEN; ANDREWS; PLISETSKAYA, 1987; PLISETSKAYA; MOMMSEN, 1996; PLISETSKAYA, 1993). Furthermore, the zebrafish GLP-1R has been cloned and found to bind with a similar affinity to both zebrafish and human GLP-1 (MOJSOV, 2000). Even more relevant to our study, the physiological roles of GLP-1 in the brain of teleost fish seem to be analogous to those described for mammals (MOJSOV, 2000; MOMMSEN; MOJSOV, 1998; PLISETSKAYA; MOMMSEN, 1996; SILVERSTEIN et al., 2001). We thus aimed to investigate the effects of liraglutide on stress-related outcomes relevant to anxiety and mood disorders on behavioral and neurochemical assays in zebrafish. More specifically, we examined the effects of liraglutide on behavior and oxidative status parameters altered by acute and chronic stress in adult zebrafish.

## 2. MATERIAL AND METHODS

### 2.1. Animals and housing standards

A total of 474 adult zebrafish of the outbred wild-type short-fin phenotype (*Danio rerio*, F. Hamilton 1822, 6-month-old, 3-4 cm long) of both sexes (~50:50 male: female ratio) were obtained from a commercial supplier (Delphis, RS, Brazil). The animals were maintained in our facility (Altamar, SP, Brazil) according to established protocols (WESTERFIELD, 2007). After the quarantine period (30 days), animals were transferred to 16-L home tanks (40 x 20 x 24 cm) with non-chlorinated water kept under constant mechanical, biological and chemical filtration, where they acclimated for another five days before testing. Fish were fed three times a day with brine shrimp (*Artemia salina*) and commercial flake fish food (Poytara^®^, SP, Brazil). The Ethics Committee of the Universidade Federal do Rio Grande do Sul approved all protocols (#32485/2017). The manipulation and animal care were conducted according to the National Institute of Health Guide for Care and Use of Laboratory Animals and aimed to minimize the discomfort, suffering, and the number of animals used.

### 2.2. Drug administration

Liraglutide (LIR, Victoza^®^, Novo Nordisk, Denmark) and 0.9% sodium chloride solution (saline, ADV Farma, SP, Brazil) were acquired from a commercial supplier. Tricaine (MS-222, CAS #886-86-2) was acquired from Sigma-Aldrich. Drug solutions were prepared daily based on previous studies and a pilot study (unpublished data). Intraperitoneal (i.p.) injections were applied using a Hamilton Microliter™ Syringe (701N 10 μL SYR 26s/2″/2) x Epidurakatheter 0.45 x 0.85 mm (Perifix^®^-Katheter, Braun, Germany) x Gingival Needle 30G/0.3 x 21 mm (GN injecta, SP, Brazil). The injection volume was 1 μL/100 mg of animal weight. The animals of the control group received the same volume of saline solution (0.9% NaCl). The animals were anesthetized by immersion in a solution of tricaine (300 mg/L) until loss of motor coordination and reduced respiratory rate. The anesthetized fish were gently placed in a sponge soaked in water submerged in a petri dish, with the abdomen facing up and the fish's head positioned on the sponge hinge. The needle was inserted parallel to the spine in the abdomen’s midline posterior to the pectoral fins. This procedure was conducted in approximately 10 seconds. For acute experiments, the time between injection and the behavioral tests was 120 minutes. For chronic experiments, behavioral tests took place 24 hours after the last injection.

### 2.3. Study design

The experimental design is shown in Figure 1. In all experiments, fish were randomly allocated into the experimental groups using a computerized random number generator (www.random.org). Behavioral and biochemical assays were conducted and analyzed by investigators blind to the experimental groups (each tank was coded by a researcher who did not participate in the experiments). The codes of experimental groups were revealed only during data analysis. Each experimental group originated from two identical home tanks, and no tank effects were observed in the analysis, so data were pooled together.

**Fig. 1.**
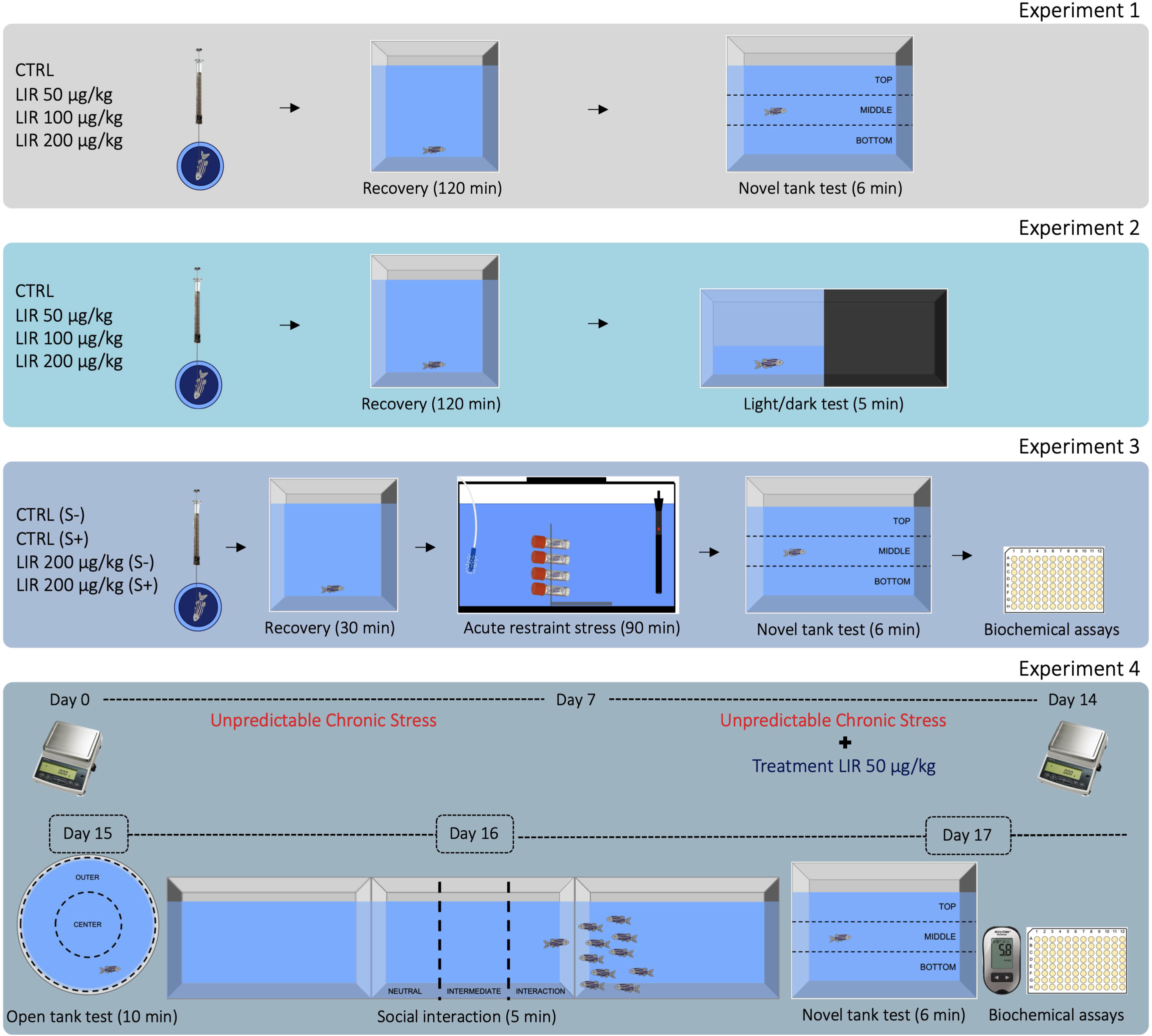
Outline of the study design.

#### Experiments 1 and 2

The animals were anesthetized and injected with either saline (CTRL) or liraglutide (LIR; 50, 100, or 200 μg/kg). After injection, fish were placed into recovery tanks (18 x 18 x 18 cm) for 120 minutes. Following recovery, fish were gently transferred to the novel tank test (NTT) in experiment 1 or to the light/dark test (LDT) in experiment 2 (MOCELIN et al., 2015; REIS et al., 2019).

#### Experiment 3

The acute restraint stress (ARS) protocol has been detailed in previous studies (DAL SANTO et al., 2014; GHISLENI et al., 2012; PIATO et al., 2011b; REIS et al., 2019). Briefly, fish were anesthetized and administered saline (CTRL) or 200 μg/kg LIR (this dose was chosen based on the anxiolytic-like effect observed in the LDT). After injection, fish were placed into recovery tanks (18 x 18 x 18 cm) for 30 min. Then, fish were allocated to stressed (ARS) or non-stressed groups. Fish in the stressed group were gently placed into 1.8-mL cryogenic tubes (Corning^®^) with openings at both ends (to allow adequate water circulation) in a 16-L tank for 90 min. Non-stressed groups were transferred to a 16-L tank for 90 min. The animals were then individually transferred to the NTT, and behavioral parameters were quantified. After the NTT, the fish were cryoeuthanized and the brain dissected and homogenized for oxidative status biochemical assays.

#### Experiment 4

The unpredictable chronic stress (UCS) protocol was carried out based on previous studies by our group (MARCON et al., 2019; MOCELIN et al., 2019b; PIATO et al., 2011a). Schedule and stressors are detailed in the supplementary material (Table S1). Initially, fish were divided into control (non-stressed group, S-) and UCS (stressed group, S+). After seven days, the experimental groups were subdivided into control (0.9% saline) and 50 μg/kg LIR (this dose was chosen because it did not alter behavior and metabolic parameters after daily injections for 7 days (Fig. S1). The animals were anesthetized daily and injected at 08:00 a.m., and then returned to the housing tanks. After UCS protocol, fish were submitted to the open tank test (OTT), social interaction (SI), and NTT, performed respectively on the 15^th^, 16^th^, and 17^th^ day. Immediately after the NTT, fish were anesthetized by rapid cooling (immersion in water at 2–4 °C until unable to swim and cessation of opercular movement); blood was collected from the tail, and the brain dissected and homogenized for biochemical assays.

### 2.4. Behavioral tests

The novel tank test (NTT) was carried out as described previously (MARCON et al., 2019; MOCELIN et al., 2015, 2019a; REIS et al., 2019; VALADAS et al., 2019). Animals were individually placed in the NTT apparatus and recorded for 6 min. The software ANY-maze™ (Stoelting Co., USA) was used to virtually divide the tank into three equal horizontal zones and track the movement of the animals. The parameters quantified were total distance traveled (m), number of crossings (transitions between the zones of the tank), number of entries, and time spent in the top zone of the tank (s).

The light/dark test (LDT) was performed as previously described by our group (MOCELIN et al., 2015; PANCOTTO et al., 2018; REIS et al., 2019). Fish were individually transferred to the white compartment of the apparatus and video recorded for 5 min. The software BORIS^®^ (Università Degli Studi di Torino, Italy) was used to quantify the time spent in the lit side (s) and the number of crossings.

The open tank test (OTT) was conducted as previously described (BENVENUTTI et al., 2020). Fish were individually placed in the center of the circular arena made of opaque white plastic (24 cm diameter, 8 cm walls), with a water depth of 3 cm, recorded for 10 min, and then analyzed using the ANY-maze™ software. The arena was virtually divided into the center (inner circle, 8 cm in diameter) and outer zones. The parameters quantified were total distance traveled (m), time immobile (s), absolute turn angle (°), and time spent in the center area of the tank (s).

The social interaction test (SI) was conducted as described previously (BENVENUTTI et al., 2020). Animals were placed for 7 min in a tank flanked by an empty tank (neutral stimulus) and a tank containing 10 zebrafish (social stimulus). Animals were habituated to the apparatus for 2 min and then analyzed for 5 min. The position of the social stimulus was counterbalanced between right or left throughout tests. The test apparatus was virtually divided into three vertical areas (interaction, middle and neutral); behavior was quantified using ANY-maze™ software. The following measures were quantified: total distance traveled, time spent in the interaction zone (s), interaction time of the fish face to face with the stimulus (s), and the number of interactions.

### 2.5. Biochemical assays

For the biochemical analyses, each sample was composed of 4 dissected brains pooled and gently homogenized in 600 μL phosphate-buffered saline (PBS, pH 7.4, Sigma-Aldrich) using a disposable tissue grinder pestle in 1.5-mL microcentrifuge tubes. Homogenates were centrifuged at 10,000 *g* at 4 °C in a cooling centrifuge, and the supernatant packed in microtubes was used for experimental assays (MOCELIN et al., 2019a). Protein was quantified according to the Coomassie blue method (BRADFORD, 1976) using bovine serum albumin (Sigma-Aldrich) as a standard. The following parameters were quantified for the oxidative status assessment: lipid peroxidation (thiobarbituric acid reactive species – TBARS) and non-protein thiols (NPSH). All biochemical measures were performed in triplicate. Details of each procedure were described in previous studies (MARCON et al., 2019; MOCELIN et al., 2019a; REIS et al., 2019; VALADAS et al., 2019). The detailed protocols for tissue preparation and biochemical assays are also available at protocols.io (10.17504/protocols.io.bjkdkks6; 10.17504/protocols.io.bjnfkmbn; 10.17504/protocols.io.bjrkkm4w; 10.17504/protocols.io.bjp8kmrw).

For the quantification of blood glucose levels, fish were anesthetized by rapid cooling, and a cut was made on the tail; the blood was collected directly on the test strips inserted in the glucometer (Accu-Chek^®^ Performa, Roche, Germany). After blood collection, the animal was euthanized by decapitation for brain tissue sampling.

### 2.6. Statistical analysis

Normality and homogeneity of variances were confirmed for all data sets using D’Agostino-Pearson and Levene tests, respectively. Results were analyzed by one, two, or three-way ANOVA followed by Tukey as applicable. For behavioral data, the outliers were identified based on distance travelled using the ROUT statistical test (GraphPad^®^ software) and were removed from the analyses. The tank and sex effects were tested in all comparisons; data were pooled when no effect was observed. Data are expressed as mean ± standard error of the mean (S.E.M). The level of significance was set at p<0.05.

## 3. RESULTS

Figure 2 shows the acute effects of liraglutide on behavioral parameters in the NTT. No significant effects of liraglutide were observed after acute treatment.

**Fig. 2.**
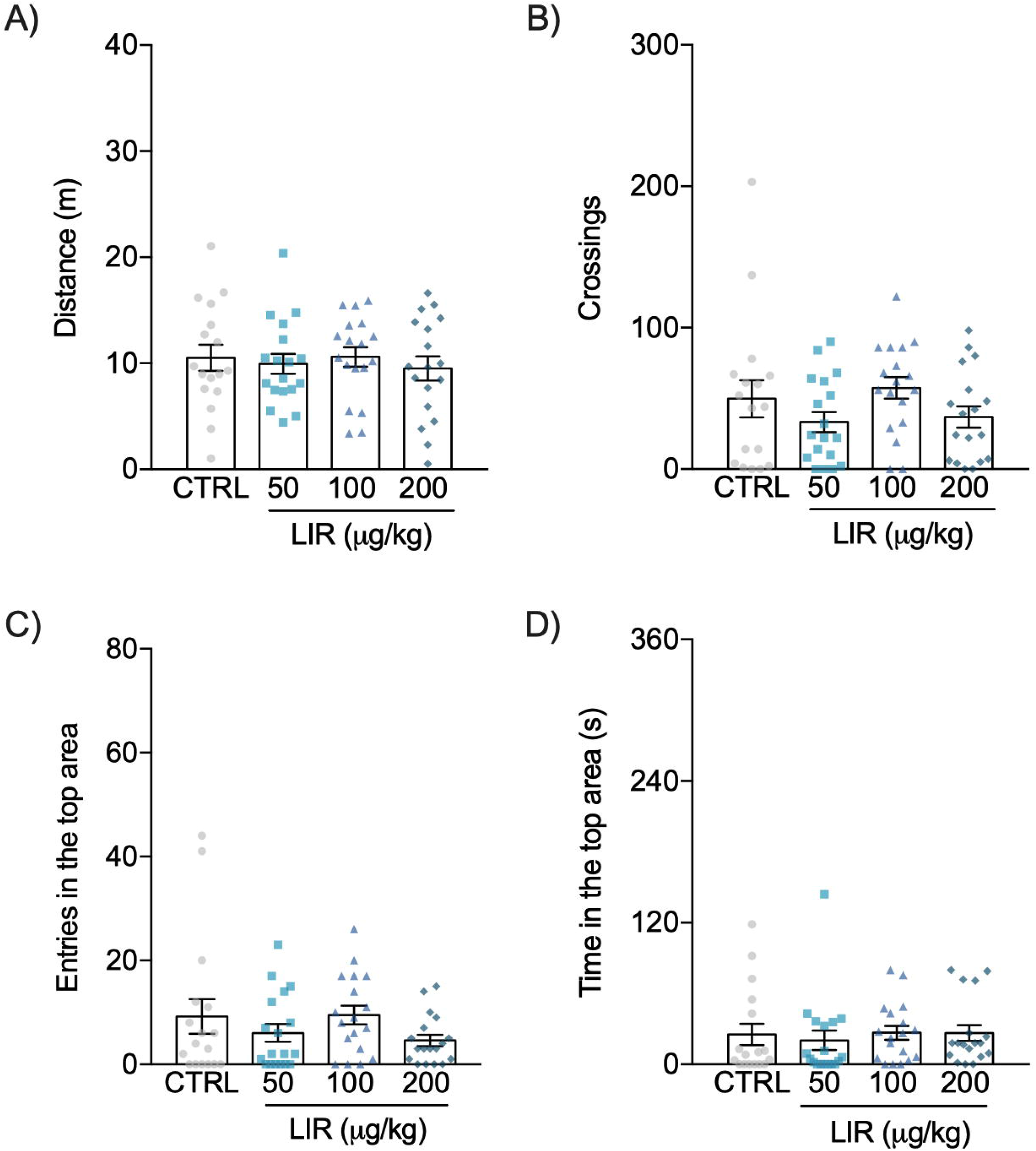
Effects of liraglutide (LIR; 50, 100, and 200 μg/kg) on distance traveled (A), number of crossings (B), entries (C), and time in the top area (D) in the novel tank test. Data are expressed as mean ± S.E.M. One-way ANOVA/Tukey. n=17-18.

Figure 3 shows the acute effects of liraglutide on behavioral parameters in the LDT. For time on the lit side, two-way ANOVA revealed an interaction between sex and treatment factors. One-way ANOVA followed by Tukey’s post hoc analysis restricted by sex, revealed that male zebrafish administered the highest liraglutide dose spent more time in the lit side than controls (p<0.05, Fig. 3A). No significant effects were observed in the number of crossings (Fig. 3B).

**Fig. 3.**
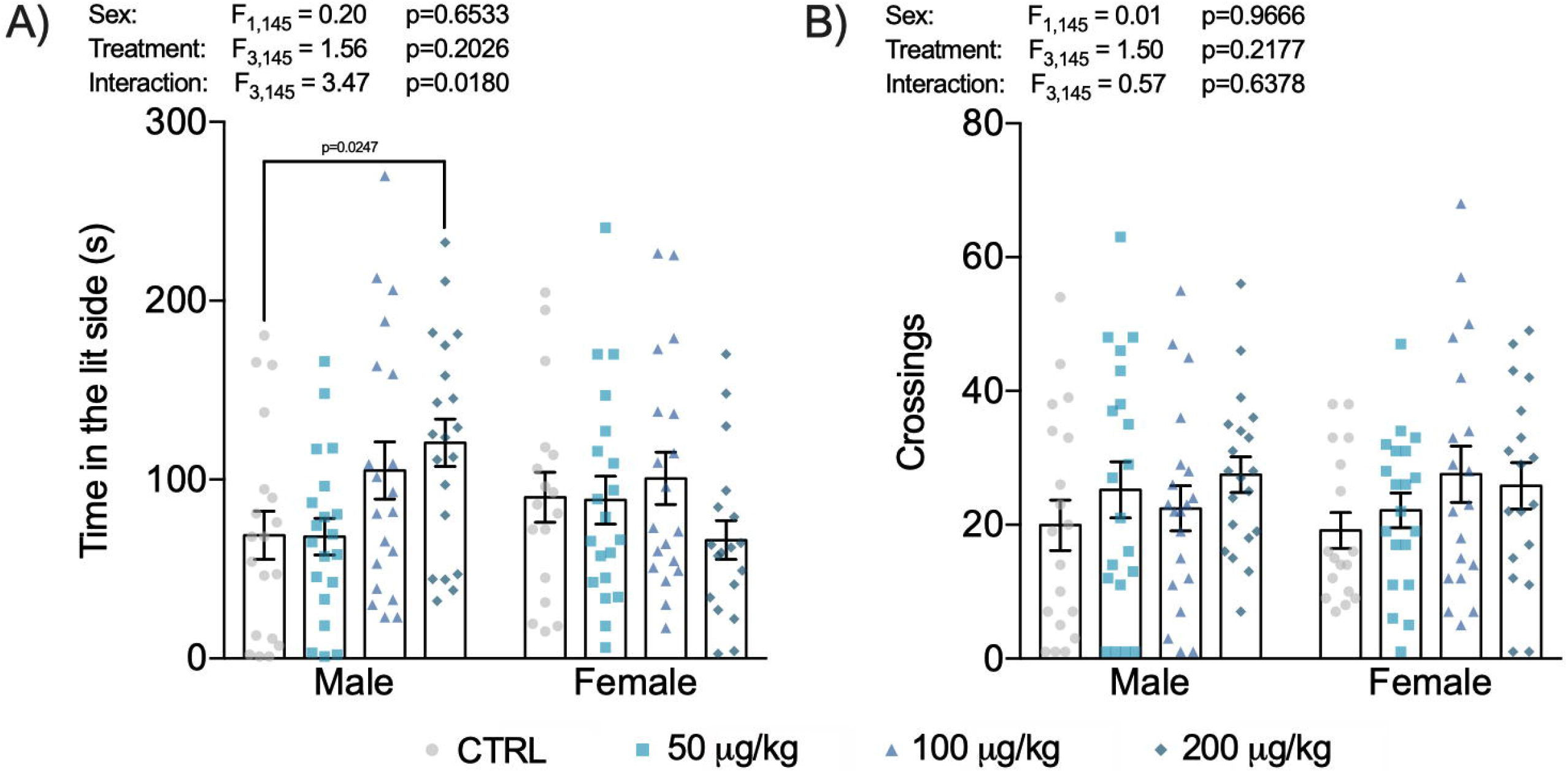
Effects of liraglutide (LIR; 50, 100, and 200 μg/kg) on time in the lit side (A) and the number of crossings (B) in the light/dark test. Data are expressed as mean ± S.E.M. Two-way ANOVA/Tukey. *p<0.05 x CTRL male. n=17-20.

The effects of liraglutide on behavioral and oxidative status parameters in zebrafish submitted to ARS and tested in the NTT are presented in Figure 4. Two-way ANOVA revealed a main effect of stress on the total distance traveled (Fig. 4A), number of crossings (Fig. 4B), and the time spent in the top area (Fig. 4D). Liraglutide did not prevent the effects of acute stress on these behavioral parameters. ARS also increased lipid peroxidation, and liraglutide was able to prevent this effect (Fig. 4E). Liraglutide did not alter the behavioral and oxidative status parameters in non-stressed animals. No effects were observed for NPSH levels.

**Fig. 4.**
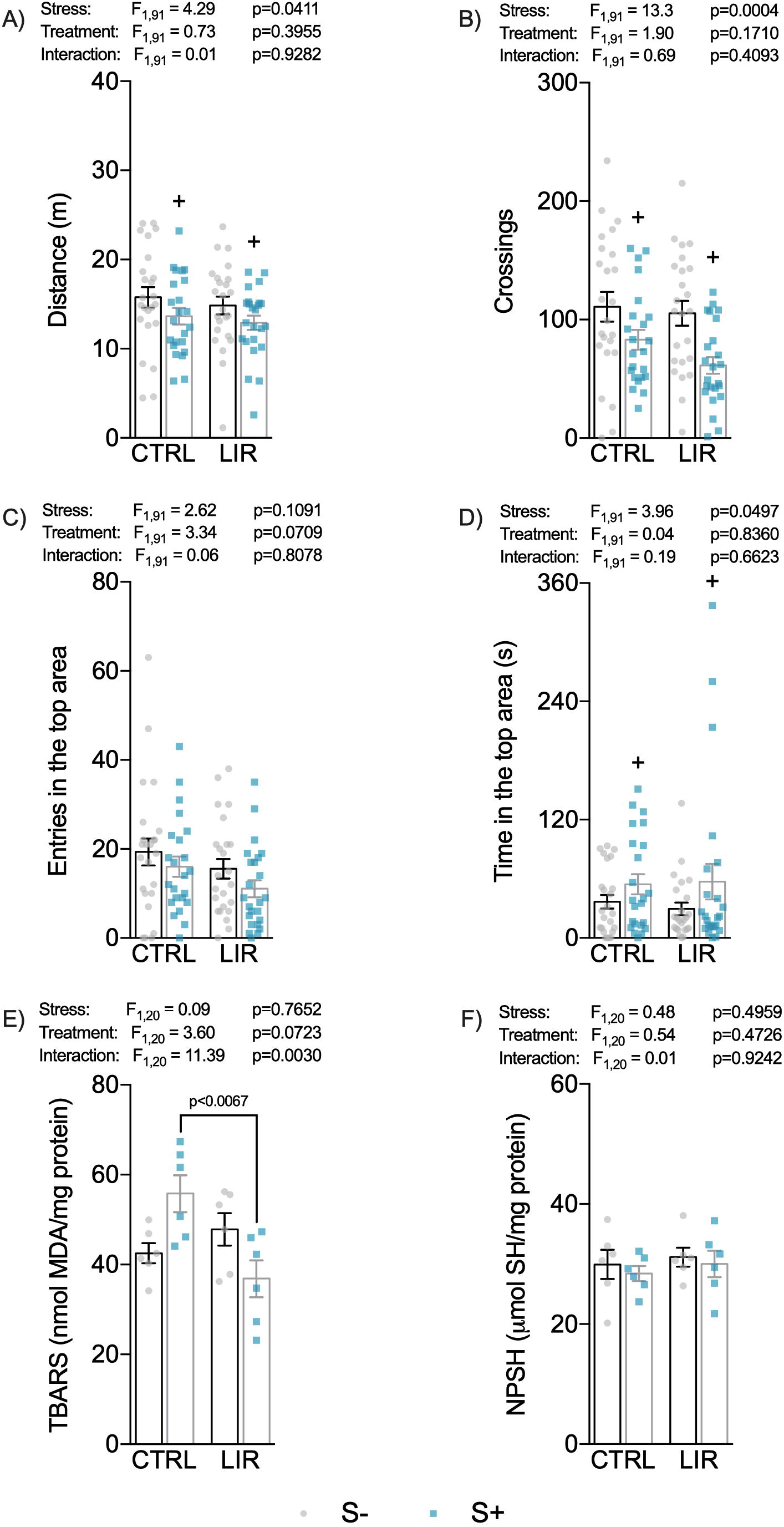
Effects of liraglutide (LIR; 200 μg/kg) on distance traveled (A), number of crossings (B), entries (C) and time (D) in the top area, lipid peroxidation (E), and non-protein thiols (F) in the acute restraint stress. CTRL, control; LIR, liraglutide; S-, non-stressed; S+, stressed. Data are expressed as mean ± S.E.M. Two-way ANOVA/Tukey. n=24 and n=6 for behavioral and biochemical analyses, respectively.

Figure 5 shows the influence of liraglutide on behavioral outcomes in zebrafish submitted to UCS. Two-way ANOVA revealed a main effect of stress on time immobile for the OTT and interaction between both factors on total distance, time immobile, and absolute turn angle. Post hoc analysis revealed that animals exposed to UCS showed a tendency for lower distance traveled (p= 0.0643), and they spent more time immobile. Liraglutide blocked these effects (Fig. 5A). No significant effects were observed in SI test (Fig. 5B). In the NTT, as expected, stress-exposed animals spent less time in the top area. Liraglutide was devoid of effects in the NTT (Fig. 5C).

**Fig. 5.**
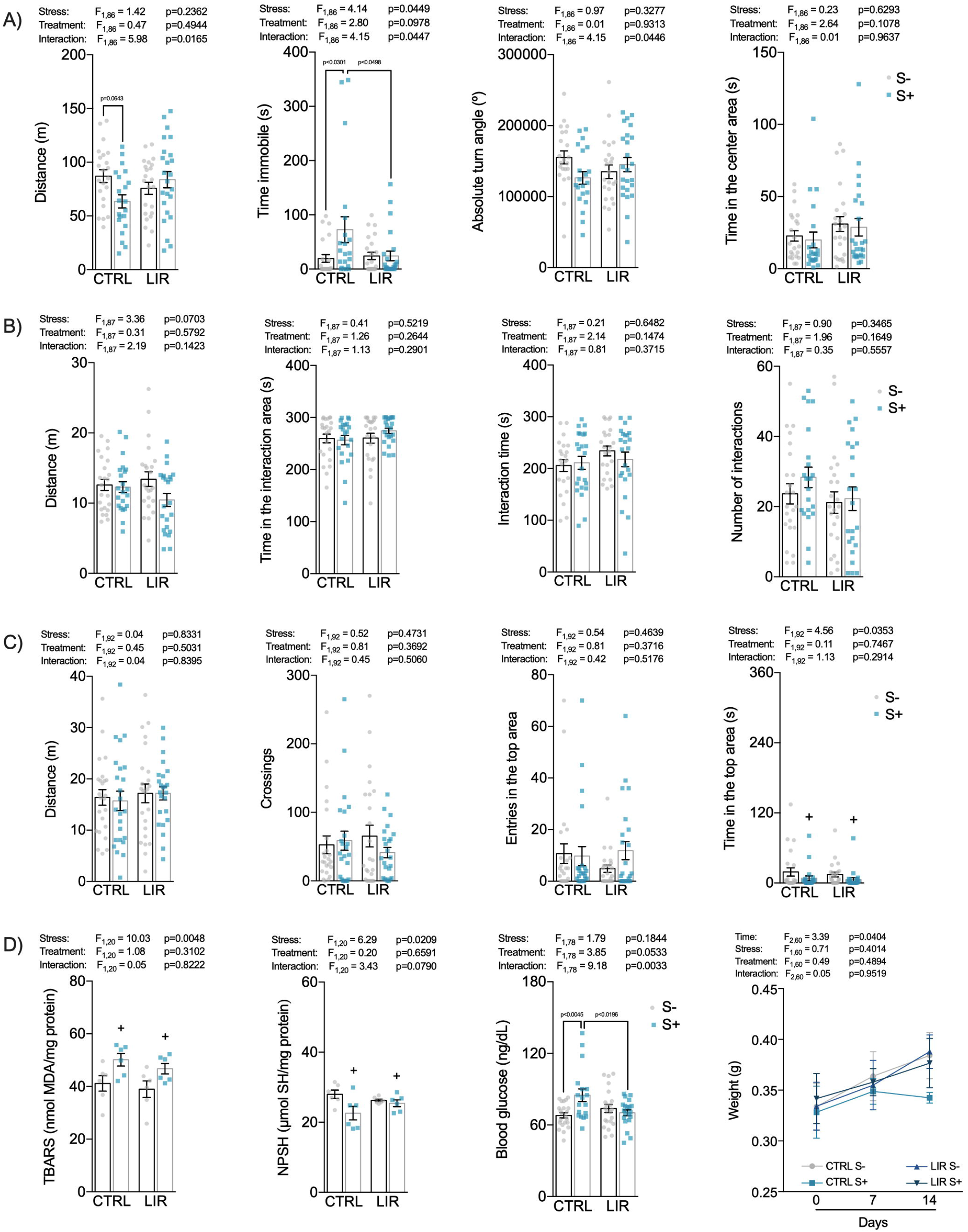
Effects of liraglutide (LIR; 50 μg/kg) in zebrafish submitted to the unpredictable chronic stress in the open tank test (Fig. 5A, day 15); in the social interaction test (Fig. 5B, day 16); in the novel tank test (Fig. 5C, day 17); in the biochemical parameters, blood glucose and weight (Fig. 5D, day 17). CTRL, control; LIR, liraglutide; S-, non-stressed; S+, stressed. Data are expressed as mean ± S.E.M. Two-way ANOVA/Tukey. n=21-24 (A); n=22-23 (B); n=24 (C); n=5-6 (D). Three-way ANOVA/Tukey was used to analyze bodyweight. n=6.

In the biochemical parameters (Fig. 5D), chronic stress increased lipid peroxidation and decreased antioxidant defenses (NPSH levels). Liraglutide was unable to block these alterations. There was an increase in blood glucose levels in stressed animals, which was blocked by treatment with liraglutide. Despite the tendency of liraglutide to block the effects of UCS on animals’ weight gain, no significant effects were observed except for the main effect of time.

## 4. DISCUSSION

In this work, we demonstrated the behavioral and biochemical effects of liraglutide administration to adult zebrafish. Initially, liraglutide was evaluated in two protocols (NTT and LDT) after a single administration to determine the effects of this drug on locomotor, exploratory, and anxiety parameters. Finally, considering the comorbidity between diabetes and stress-related disorders such as anxiety and depression, we studied the effects of liraglutide in both acute and chronic stress models.

The NTT is a vertical exploratory test in which bottom-dwelling is analog to the thigmotaxis displayed by rodents in the open-field test (OFT) (LEVIN; BENCAN; CERUTTI, 2007; MOCELIN et al., 2015). Anxiolytic drugs such as buspirone, fluoxetine, and diazepam increase the time zebrafish spend in the top area in the NTT (BENCAN; SLEDGE; LEVIN, 2009; EGAN et al., 2009; GEBAUER et al., 2011; MOCELIN et al., 2015; REIS et al., 2019). In the LDT, anxiolytic drugs such as clonazepam, bromazepam, diazepam, and buspirone, increase the time zebrafish spend in the lit side of the apparatus (GEBAUER et al., 2011; MOCELIN et al., 2015; REIS et al., 2019). Here, we found that acute treatment with liraglutide did not affect the locomotor and exploratory behaviors in zebrafish, assessed by the total distance traveled and the number of crossings between areas in the NTT. The same was observed when animals were chronically exposed to liraglutide for seven days (Fig. S1). In the LDT, the highest dose of liraglutide (200 μg/kg) induced anxiolytic-like behavior, i.e., increased the time spent in the lit side of the apparatus. This anxiolytic-like behavior in the LDT was observed only in male zebrafish. The sex-dependent effect of liraglutide needs further investigation. Previous studies in zebrafish have shown that males are generally bolder in social interactions than females (ARIYOMO; CARTER; WATT, 2013; DAHLBOM et al., 2012; MORETZ; MARTINS; ROBISON, 2007), exploring more new environments (REOLON et al., 2018), and showing greater aggressiveness and cortisol levels when exposed to chronic stress (RAMBO et al., 2017). However, females have better memory acquisition in some cognitive tests such as inhibitory avoidance (REOLON et al., 2018), but not in others such as in the plus-maze (SISON; GERLAI, 2010). Some studies have shown there are sex-dependent responses when animals are exposed to drugs of abuse such as cocaine, the effects of the withdrawal being more intense and persistent in males than in females, but the related mechanism of action remains to be investigated (LÓPEZ PATIÑO et al., 2008). In general, studies showing differences in pharmacological responses between males and females in this species are scarce and warrants further investigation.

There are many acute stress protocols established for zebrafish (ABREU et al., 2014; BLAZINA; VIANNA; LARA, 2013; PIATO et al., 2011b). For example, restraining zebrafish in a small tube induces a consistent anxiety-like behavior, decreasing the time and entries in the top area of the NTT apparatus. Despite the robust effects on anxiety outcomes, the effects of the ARS on locomotor parameters are heterogeneous, increasing (GHISLENI et al., 2012), decreasing (REIS et al., 2019), and sometimes not affecting (ASSAD et al., 2020; PIATO et al., 2011b) locomotion. These contradictory results concerning the effects of ARS on zebrafish locomotion could be related to the different protocols used (restraint time, for example) in the different studies and/or methods used for quantifying the behavior. Here, 90 min of ARS decreased the locomotor parameters and induced an apparent anxiogenic behavior, whereas liraglutide did not prevent these phenotypes. Moreover, ARS increased lipid peroxidation. This result is in agreement with our previous study (DAL SANTO et al., 2014). Liraglutide presented neuroprotective effects in our study, blocking the lipoperoxidation induced by acute stress. The effects of GLP-1R agonists in animal models of anxiety are conflicting. For example, intracerebroventricularly-injected GLP-1 induced anxiogenic effects in the elevated plus-maze test in rats (GULEC; ISBIL-BUYUKCOSKUN; KAHVECI, 2010). Moreover, liraglutide was anxiogenic in the OFT in rats (KAMBLE et al., 2016). However, exenatide showed anxiolytic- and antidepressant-like effects in a diabetic mouse model (KOMSUOGLU CELIKYURT et al., 2014) and liraglutide induced anxiolytic effects in elevated plus-maze in rats (SHARMA et al., 2015). In a mice study, both exenatide and liraglutide did not affect anxiety levels in the light/dark test, but both decreased distance and rearing (KRASS et al., 2012). Therefore, the *per se* effects of GLP-1R agonists are very conflicting, and it can be explained, at least in part, by the different doses used in these studies.

Mental disorders such as depression and anxiety present complex neurobiology, and the mechanisms related to these conditions are not entirely understood. The gut-brain axis and neuropeptides like GLP-1 are rising as potential key regulators of behavior (POZZI et al., 2019). Animal models of chronic stress have been used to recapitulate the phenotypes observed in patients with mood disorders (WILLNER, 2005, 2017). Specifically, in zebrafish, the UCS induces behavioral, neurochemical, and molecular alterations that resemble the clinical presentation of patients with anxiety or depressive disorders. Antidepressants and anxiolytics have been previously shown to block the effects of chronic stress in zebrafish (MARCON et al., 2016; SONG et al., 2017). Moreover, drugs with antioxidant and neuroprotective mechanisms have also shown protective effects in this model (MARCON et al., 2019; MOCELIN et al., 2019b). Here, we submitted zebrafish to UCS for two weeks. After this period, zebrafish behavior was assessed in three tests: OTT, SI, and NTT. We extended the behavioral outcomes reported in previous studies from our group to expand the characterization of the impact of UCS in zebrafish. In the OTT, UCS decreased the distance and absolute turn angle and increased the immobility time. Liraglutide was able to block these effects. As expected, in the NTT, the UCS decreased the time in the top area. In the social interaction test, however, no significant effects were observed. Our results replicate and reinforce previous studies showing that UCS induces anxiety-like behavior in the NTT (MARCON et al., 2016, 2019; MOCELIN et al., 2019b; PIATO et al., 2011a). Liraglutide did not prevent the effects of UCS on those last two behavior tests. UCS also increased lipid peroxidation and decreased the levels of an important antioxidative defense in the brain. Previous studies reported that UCS promoted an imbalance in oxidative status, increased lipoperoxidation, and decreased antioxidant defenses (MARCON et al., 2018, 2019; MOCELIN et al., 2019b). Although liraglutide counteracted lipid peroxidation induced by an acute stressor, it did block the neurochemical effects of chronic stress. In rodents, there is some evidence regarding the positive effects of GLP-1R agonists in chronic stress. For example, liraglutide diminished depressive- and anxiety-like phenotypes induced by chronic administration of corticosterone. This effect can be related to modulation of the hypothalamus-pituitary-adrenal (HPA) axis (WEINA et al., 2018). In contrast, another study has shown no benefit of using GLP-1R agonists in models of depression/anxiety. Fourteen days of treatment with exenatide or liraglutide did not affect the anxiety level in LDT nor induced an antidepressant-like effect in the forced swimming test in mice (KRASS et al., 2015). In the same study, chronic treatment with liraglutide did not affect depression-related behaviors in Flinders Sensitive Line rats, a genetic model of depression. These divergent results can be explained, at least in part, by the doses and the administration route used in the studies. Despite these inconsistencies, in our study, liraglutide prevented some, but not all, behaviors induced by UCS.

Here, UCS increased glycemia and decreased weight gain. Liraglutide prevented the effects of UCS on glycemia without effects on zebrafish weight. In all doses, liraglutide per se did not alter the glycemia (see Fig. S1). In higher doses, the drug affected the weight gain after 7-days treatment (Fig. S1). Although liraglutide induces a clear meal-dependent hypoglycemic effect in mammals through interaction with GLP-1R, the peripheric effects of GLP-1R agonists in the teleost fish are distinct (MOJSOV, 2000). GLP-1 has a glucagon-like activity in teleost fish, increasing glycemia through hepatic glycogenolysis and gluconeogenesis, an antagonistic response to insulin (MOMMSEN; ANDREWS; PLISETSKAYA, 1987; PLISETSKAYA; MOMMSEN, 1996; YEUNG et al., 2002). Despite this divergent response between mammals and fish, the central effects are conserved, as GLP-1 controls feeding behavior in the same manner in both groups (SILVERSTEIN et al., 2001; TURTON et al., 1996). Despite the hyperglycemic effects of GLP1-R activation in zebrafish, here liraglutide prevented the increase in circulating glucose induced by UCS. Although this may be an unexpected result, the effects of GLP-1R agonists in zebrafish are poorly understood.

The high comorbidity between mental disorders and diabetes is explained by a myriad of interactions including central and peripherical factors as a bidirectional association that results in not only alterations in the glucose and insulin metabolism but also oxidative damage and inflammation (CARTER; SWARDFAGER, 2016; GRIGOLON et al., 2019; NEFS et al., 2012, 2019). The role of GLP-1 in the stress response is recent (GHOSAL; MYERS; HERMAN, 2013). Few studies have investigated GLP-1R agonists’ effects in acute/chronic stress, even in mice and rats. In zebrafish, we suggest liraglutide improves the global stress response, and these effects can be explained, at least in part, by its antioxidant effects, but the mechanisms need to be better studied.

## 5. CONCLUSION

Liraglutide is part of a new generation of drugs approved for glycemic control in diabetic patients. Recent findings suggested GLP-1R agonists may be effective in treating mental disorders, but only a few studies assessed their effects in stress-related mental disorders. Our study adds to a growing body of literature demonstrating GLP-1R agonists’ role in modulating behavior and homeostasis after acute or chronic stress. Whereas diabetic patients are prone to developing mental disorders, a drug able to treat both conditions would reduce costs, adverse effects, and nonadherence to treatment. Therefore, GLP-1R agonists are potential candidates to be repurposed for mental disorders; however, more studies are necessary to elucidate the mechanisms of liraglutide in preclinical and clinical studies.

## Supporting information

Supplemental_Bertelli et al., 2021

## AUTHOR CONTRIBUTION

Conceived and designed the experiments: PB, RM, AR, RG, AP; Performed the experiments: PB, RM, MM, AS; Analyzed the data: PB, RM, AS, AH, AP; Contributed reagents/materials/analysis tools: AP; Wrote the paper and prepared figures: PB, RM, AH, AP; Authored or reviewed drafts of the paper: all authors; Approved the final draft: all authors.

## ACKNOWLEDGMENTS

The authors extend their sincere thanks to the LAPCOM team for their contribution.

## ETHICAL STATEMENT

The Ethics Committee of the Federal University of Rio Grande do Sul approved all protocols (process number # 32485). The manipulation and animal care were conducted according to the National Institute of Health Guide for Care and Use of Laboratory Animals and aimed to minimize the discomfort, suffering, and the number of animals used.

