## Supplemental_Bertelli et al., 2021 for "Anti-stress effects of the glucagon-like peptide-1 receptor agonist liraglutide in zebrafish"

*Correspondence to:

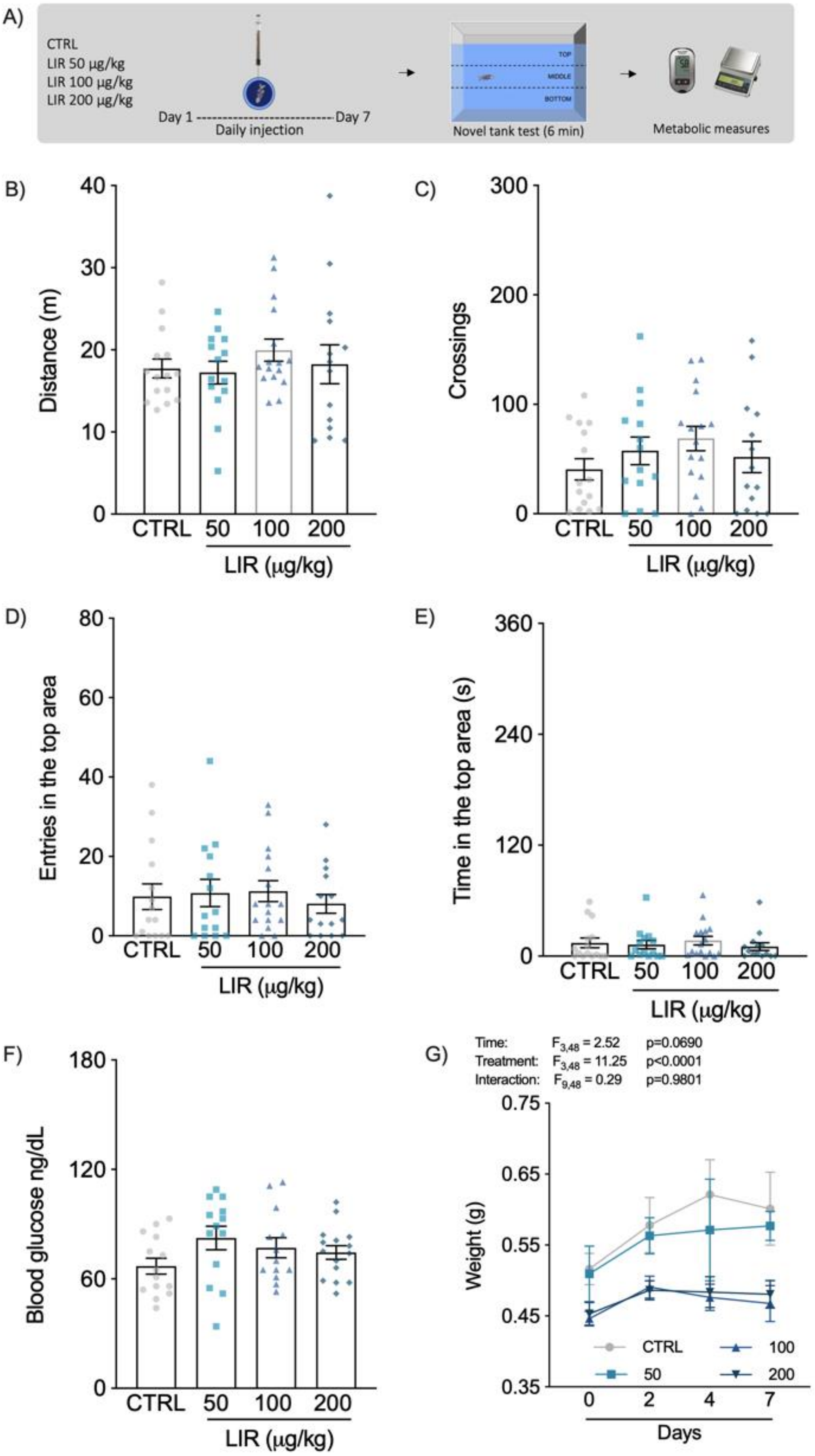


**Fig. S1.** Experimental design (S1A). Effects of liraglutide (LIR; 50, 100, and 200 µg/kg) on behavior (distance traveled [S1B], number of crossings [S1C], entries [S1D] and time [S1E] in the top area in the novel tank test). Figures S1F and S1G show the effects of liraglutide (50, 100, and 200 µg/kg) on blood glucose and weight. Data are expressed as mean ± S.E.M. One-way ANOVA/Tukey (n=14-16). Bodyweight was analyzed by two-way ANOVA/Tukey (n=4).

Figure S1 shows the chronic effects of liraglutide on behavioral, blood glucose, and weight parameters. Figure S1A represents experimental design. One-way ANOVA followed by Tukey post hoc analysis showed no significant effects of liraglutide treatment on the behavioral parameters in the NTT (Fig. S1B-D). Regarding blood glucose, one-way ANOVA did not reveal any effects of liraglutide after seven days of treatment (Fig. S1F). Two-way ANOVA revealed a main effect of treatment, where animals treated with the highest doses of liraglutide (100 and 200 μg/kg) had lower bodyweights as compared to controls (Fig. S1G).

| DAY | 1 | 2 | 3 | 4 | 5 | 6 | 7 |
|---|---|---|---|---|---|---|---|
| MORNING | 8h15 | 12h20 | 11h30 | 9h30 | 11h40 | 12h30 | 8h00 |
| | tank change, three consecutive times with 30 min interval | crowding in a 250-mL beaker (50 min) | heating tank water to 18˚C (30 min) | low water level on housing tanks until dorsal body wall was exposed (5 min) | cooling tank water to 18˚C (30 min) | low water level on housing tanks until dorsal body wall was exposed (5 min) | crowding in a 250-mL beaker (50 min) |
| AFTERNOON | 12h55 | 16h15 | 15h30 | 16h30 | 16h00 | 15h30 | 15h30 |
| | cooling tank water to 18˚C (30 min) | Chasing with a net (8 min) | crowding in a 250-mL beaker (50 min) | tank change, three consecutive times with 30 min interval | crowding in a 250-mL beaker (50 min) | heating tank water to 18˚C (30 min) | Chasing with a net (8 min) |
| DAY | 8 | 9 | 10 | 11 | 12 | 13 | 14 |
| MORNING | 12h00 | 8h00 | 10h30 | 9h00 | 9h30 | 12h00 | 8h45 |
| | tank change, three consecutive times with 30 min interval | crowding in a 250-mL beaker (50 min) | low water level on housing tanks until dorsal body wall was exposed (5 min) | heating tank water to 18˚C (30 min) | tank change, three consecutive times with 30 min interval | low water level on housing tanks until dorsal body wall was exposed (5 min) | crowding in a 250-mL beaker (50 min) |
| AFTERNOON | 17h30 | 12h30 | 15h30 | 16h00 | 13h30 | 17h10 | 16h30 |
| | cooling tank water to 18˚C (30 min) | Chasing with a net (8 min) | crowding in a 250-mL beaker (50 min) | tank change, three consecutive times with 30 min interval | heating tank water to 18˚C (30 min) | crowding in a 250-mL beaker (50 min) | Chasing with a net (8 min) |

**Table. S1.** Schedule and stressors of the unpredictable chronic stress protocol.
